# Transferrin Receptor 1-targeted polymersomes therapy for Colorectal Cancer

**DOI:** 10.1101/2025.02.24.639871

**Authors:** Ariana Pina, Elisa Mastrantuono, Marta Silva, Valentino Barbieri, José Muñoz-López, Giuseppe Battaglia, Luís Graça, Diana Matias

## Abstract

Colorectal cancer (CRC) is a major global health concern, ranking among the most common cancers and the second leading cause of cancer-related deaths. The high mortality associated with CRC is attributed mainly to difficulties in early detection and the lack of effective targeted therapies. The Transferrin receptor 1 (TfR1) is particularly attractive as a therapy target given its notable overexpression in tumor cells, particularly in CRC. This study explored the potential of a polymeric nanoparticle (POs)-based drug delivery system targeting TfR1 to improve the precision and efficacy of CRC treatment. For this study, we used three human colorectal cancer cell lines (SW480, HT-29, and HCT116), a healthy human intestinal epithelial cell line (hIECs), and a murine CRC cell line (MC38). We first engineered POs composed of poly (ethylene glycol) (PEG) and poly (lactic acid) (PLA), functionalized with the T7 peptide to enhance their specificity for TfR1-expressing cells. Targeting efficiency of these POs was assessed across all cell lines by evaluating the cellular uptake using flow cytometry. Upon establishing the optimal formulation for these NPs for TfR1-targeting, we encapsulated doxorubicin (DOX) to evaluate their therapeutic potential. Both *in vitro* and *in vivo* studies were performed to assess the efficacy of these DOX-loaded TfR1-targeted POs. *In vitro* studies demonstrated selective delivery of DOX to CRC cells, suggesting a marked reduction in off-target effects. *In vivo* studies in a murine model of CRC further supported these findings, showing that DOX-loaded TfR1-targeted POs significantly improved survival rates and reduced tumor growth compared to free DOX or PBS treatments. These results highlight the promise of TfR1-targeted POs as a precise strategy for CRC therapy, offering enhanced treatment efficacy while reducing systemic toxicity. This novel approach could lead to the development of more targeted and less harmful cancer treatments.

**Graphical abstract:** 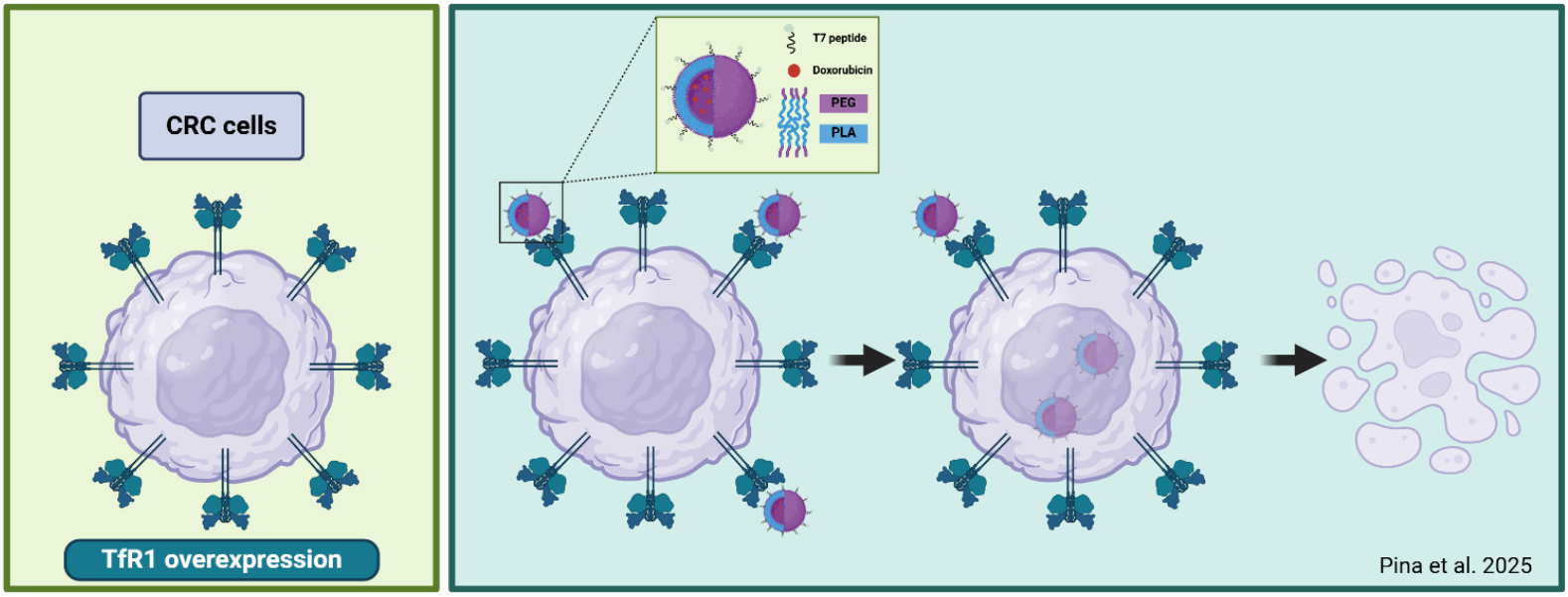

**Highlights:**

- T-7 peptide-functionalized polymersomes exhibit high binding affinity to CRC cell lines
- Doxorubicin-loaded T7-peptide polymersomes effectively suppress tumor growth and prolong survival in vivo
- Targeted Doxorubicin delivery via T7-peptide polymersomes minimizes toxicity to healthy tissues

## 1. Introduction

Colorectal cancer (CRC) remains a global health challenge, ranking as the third most prevalent cancer and the second leading cause of cancer-related deaths worldwide [1]. CRC is highly heterogeneous, both within individual tumors and across patients, with factors such as tumor location and interactions with immune cells contributing to its complex pathogenesis [2,3]. Consequently, treatment outcomes are often unpredictable, posing a challenge for conventional therapies. Standard CRC treatment typically involves surgical resection followed by chemotherapy and/or radiotherapy [4,5]. However, effectiveness depends largely on early detection, which is only achieved in approximately 40% of cases, and even when successful, recurrence rates are high, making chemotherapy become a primary option of treatment, yet it is often associated with suboptimal response rates and treatment resistance [4,6]. This resistance, along with the lack of specificity of chemotherapeutic agents, contributes to significant collateral damage to healthy tissues, manifesting as neuro-, cardio-, hepato-, and nephrotoxicity. Such side effects can exacerbate patient morbidity, reduce quality of life, and, in some cases, lead to secondary malignancies [4,7].

One of the most promising chemotherapeutic agents is doxorubicin (DOX), a well-known topoisomerase II inhibitor with high effectiveness against various malignancies, including but not limited to carcinomas, sarcomas and breast cancers [8,9]. Additionally, it is recognized for its cost-effectiveness and broad-spectrum activity in cancer treatments [10,11]. However, in the context of CRC, DOX’s efficacy is significantly limited due to several mechanisms of innate resistance, limiting its clinical application for these tumors [12].

In light of these limitations, novel strategies are needed to enhance the specificity and efficacy of CRC treatments. One effective strategy is to exploit the overexpression of receptors on tumor cells, which are often present at higher levels compared to healthy tissues, and to leverage nanotechnology for targeted drug delivery. A highly promising target in CRC is Transferrin Receptor 1 (TfR1), a type II transmembrane glycoprotein that plays a crucial role in cellular iron uptake. While TfR1 is typically expressed at low levels in healthy tissues, it is often overexpressed in cancer cells, including CRC [13]. This overexpression is strongly associated with increased cell proliferation, invasion, and metastasis, making TfR1 an attractive target for drug delivery systems, such as nanoparticles [13]. Among various polymer-based nanoparticles, specifically polymersomes (POs) stand out due to their ability to be surface-functionalized to target specific receptors, such as TfR1, without compromising their structural integrity, making them ideal for precise and efficient drug delivery [14–20].

The incorporation of FDA-approved poly(ethylene glycol) (PEG) and poly(lactic acid) (PLA) in POs enhances their biocompatibility and *in vivo* stability, further supporting their safety profile[14,15]. Additionally, this unique composition allows for customization in size, shape, and membrane properties, optimizing both drug loading and targeting efficiency for improved therapeutic outcomes [14,15].

Although TfR1 has been widely explored as therapeutic target in various cancers, its application in POs-based therapies for CRC remains largely unexplored [16]. One of the key challenges in targeting TfR1 is the competition between targeting peptides and endogenous transferrin, which significantly reduces treatment efficacy [13,16]. Overcoming this limitation is crucial to unlocking the full therapeutic potential of TfR1-targeted PO therapies in CRC.

In this study, we developed TfR1-targeted PEG-PLA POs functionalized with the T7 peptide (HAIYPRH), a ligand known for its high specificity and affinity for TfR1. These T7-functionalized POs were designed to efficiently encapsulate and deliver DOX to CRC cells. Although DOX is not a first-line treatment for CRC, it remains a potent chemotherapeutic agent across multiple cancer types. However, its clinical application in CRC is often limited by resistance mechanisms and systemic toxicity. By employing a targeted delivery strategy, we aim to enhance DOX efficacy, reduce off-target effects and bypass resistance mechanisms, thereby offering a promising therapeutic alternative for CRC patients.

## 2. Material and Methods

### 2.1. Synthesis and purification of empty POs

The PEG_45_-PLA_106_ copolymers were purchased from Creative PEGWorks. The T7 conjugated copolymers (HAIYPRH-PEG_20_-PLA_112_) were synthesized and functionalized in-house. For polymer synthesis, N3-PEG-PLA was prepared using N3-dPEG24-OH (#PEG3770, Iris Biotech) as the initiator and polymerizing LA by using ring-opening polymerization with 1,8-Diazabicyclo[5.4.0]undec-7-ene (#139009, Sigma-Aldrich) as the catalyst. Afterwards, to synthesize T7 peptide (HAIYPRH)-PEG_20_-PLA_112,_ copper(I)-catalyzed azide-alkyne cycloaddition (CuAAC) click reactions were used, with {Propargylamine}T7 peptide (GenScript) and N3-PEG_24_-PLA_112_ reacting in the presence of copper(II) sulfate (#209198, Sigma-Aldrich) as the catalyst and sodium ascorbate (#A7631, Sigma-Aldrich) as the reducing agent. PEG-PLA POs with different % molar amounts of T7 peptide were obtained using the solvent displacement method (Supp.Table1). To achieve this, PEG_45_-PLA_106_ polymer (20 mg) was dissolved in 1 mL of dimethylformamide (DMF) and vortexed to ensure dissolution. Different formulations were obtained by varying the ratio of T7 conjugated copolymers (T7-PEG_20_-PLA_112_) to the PEG_45_-PLA_106_. The solution was transferred to a syringe and 600 μL were injected into a glass vial placed on a stirring plate at 500 rpm, containing a magnet and 1600 μL of MiliQ Water. The polymer solution was injected into the MiliQ water at the steady rate of 100 μL/min. After the injection, the solutions were transferred to dialysis membranes (#Z726176, Merck) and put inside a beaker full of MiliQ water and left to stir for an hour. Then the MiliQ water was substituted by PBS and left to stir overnight. In these POs were also included a fluorophore: Cy7-PEG_20_-PLA_112_. After dialysis, samples were collected and transferred to 15 mL tubes and centrifuged at 1000 rpm for 10 minutes. These POs will be referred to as X%T7pep-POs from here onwards.

### 2.2 Characterization of POs by Dynamic Light Scattering (DLS) and transmission electron microscopy (TEM)

Hydrodynamic diameter and polydispersity (PDI) were evaluated by dynamic light scattering (DLS) using a Zetasizer Ultra. 10 3.4.4 (Malvern Instruments) with a 10 mW He-Ne Laser at 632.8 nm in a scattering angle of 173º. The POs samples were prepared by diluting 50 μL of POs into 850 μL of MiliQ water and transferred into polystyrene disposable cuvettes at 25ºC. Data was processed using a ZS XPLORER software (Malvern instruments).

For TEM, Grids composed of 100 mesh formvar, and carbon coat were glow discharged prior to the experiment and samples were diluted in a 1:5 proportion in water and sonicated for a total of 5 minutes. Grids were deposited on top of 5 μL of diluted sample for adsorption for 2 minutes and washed in distilled H_2_O 10 times. Posteriorly, the grids containing the sample were stained in 2% Uranyl Acetate in dH_2_O for 2 minutes and left to dry in Whatman filter paper. Imaging was done using a Tecnai G2 Spirit BioTWIN Transmission Electron Microscope from FEI and pictures were acquired using an Olympus-SIS Veleta CCD Camera. Stainings and acquisition were performed by the Electron Microscopy Facility at the Gulbenkian Institute for Molecular Medicine (GIMM).

### 2.3 Quantification of POs

The concentration of PEG-PLA in each formulation was quantified using ultra high-performance liquid chromatography (uHPLC) (Agilent 1260 Infinity II) with a Jupiter C18 column (Phenomenex). For PEG-PLA sample analysis, a gradient method was employed with 50% (v/v) acetonitrile (Phase A) and 50% (v/v) ultra-pure water (Phase B). The gradient progressively increased from 50% A to 100% A over 20 minutes in a linear fashion. This concentration was maintained for 25 minutes before returning to 50% A over 1 minute and holding for 6 minutes. The flow rate was set at a constant 1 mL/min. Before uHPLC analysis, POs samples were diluted in acetonitrile to facilitate polymer disassembly. POs absorbance was detected at 220 nm using a UV-Vis detector. Absorbance curves were analyzed using Chromaleon Chromatography Data System (CDS) software.

### 2.4 Synthesis and purification of POs with encapsulated DOX

The same protocol was used to create POs with encapsulated DOX (# D2975000, Merck), however, before injecting the solution into the glass vial on the stirring plate, 333 μL of DOX at a concentration of 2mg/mL were added. Furthermore, no fluorophores were added to these POS as DOX already presents fluorescence. POs were first filtered by using a crossflow filtration technique and then by size exclusion chromatography. A Size Exclusion Column (SEC) was used specifically for DOX encapsulating POs to separate them from the remaining free drug present in the samples. The samples were run through a 53 ml glass liquid chromatography column containing Sepharose 4B (#4B200-500ML, Sigma-Aldrich) as the stationary phase and PBS-endotoxin free (#TMS-012-A, Merck) as the eluent. After running through the column, POs were collected in a 96 well plate (around 200 μL per well) until PBS was clear and analyzed by Dynamic Light Scattering (DLS).

### 2.5 DOX loading efficacy and quantification in POs

To calculate the amount of encapsulated DOX in the POs, samples were diluted in a 1:5 ratio acetonitrile to water and plated in a black flat-bottom 96-well plate. A standard curve of free DOX in acetonitrile/water was also prepared, starting at a concentration of 1 mg/mL and subjected to serial dilutions in 1:2 ratio. Fluorescence was measured using a TECAN Spark microplate reader, with an excitation wavelength of 470 nm and an emission wavelength of 560 nm. The Concentration of DOX was determined by referencing the standard curve.

### 2.6 In vitro Colorectal cancer and human intestinal epithelial cultures

Colon carcinoma cell lines from human (HCT116 (#CCL-247, ATCC), HT-29 (#HT29, Creative Biogene) and SW480 (#CCL-228, ATCC) cell lines) and mouse (MC38) gently provided by (Centro de Inmunologia Molecular (CIM), Cuba) were cultured in Minimum Essential Medium Eagle (MEME) (#M4655, Sigma-Aldrich) supplemented with 10% Fetal Bovine Serum (FBS) (#10500-064, Life Technologies) and 1% Penicillin Streptomycin (P/S) (#15140-122, Life Technologies). Human Intestinal Epithelial cells (HIEC-6 cell line) (#CRL-3266, ATCC) were cultured in DMEM/F12 (#D0697-500ML, Merck) supplemented with 5% FBS, 1% Insulin-Transferrin-Selenium (#41400045, Thermo Scientific), 0.5 ng/mL of Epidermal Growth Factor (EGF) (#315-09-100 μg, PeproTech France) and 1% P/S. All cell lines were maintained in an incubator at 37ºC, 5% CO_2_ and constant humidity.

### 2.7 Evaluation of TfR1 levels in CRC and HIEC cells by immunofluorescence and Flow cytometry

Monolayers of HCT116, HT-29, HIEC-6, and MC38 cells were cultured in ibidi μ-Slide 8-well plates (#80807, Enzifarma) at a density of 20,000 cells per well, with 200 μL of their respective media. After 48 hours of incubation, the cells were washed with 1x PBS and fixed with 4% paraformaldehyde (PFA) (#P6148-1KG, Sigma-Aldrich) in PBS for 10 minutes. Subsequently, the cells were permeabilized with 0.1% Triton X-100 solution (#93443, Sigma-Aldrich) in PBS for 15 minutes at room temperature. Afterward, the cells were blocked with 5% bovine serum albumin (BSA) (#071M1561V, Sigma-Aldrich) in PBS with 0.1% Tween-20 (#P1379, Sigma-Aldrich) (PBS-T) for 1 hour, followed by overnight incubation with the primary antibody anti-TfR1 (#ab84036, Abcam, 1:200) diluted in 1% BSA in PBS-T at 4ºC. Following the primary incubation, the cells were washed with PBS and incubated with the secondary antibody anti-rabbit IgG DyLight 488 (#406404, BioLegend, 1:200) diluted in PBS. Cells were incubated for 1 hour at room temperature (RT) in a humidified, dark chamber. The cells were then washed with PBS, stained with DAPI at 0.1 μg/mL for 1 minute, washed again with PBS, and visualized using the Zeiss Cell Observer microscope (Axio Observer, ZEISS Microscopy), at the Bioimaging Platform of the GIMM. Images obtained were analyzed and processed using Fiji/ImageJ version 1.53t software.

For flow cytometry analysis, 2 x 10^5^ cells from each cell line were collected, washed twice with PBS + 2% FBS for 3 minutes at 1300 rpm, and incubated with LIVE/DEAD® Fixable Aqua Dead Cell Stain Kit, for 405 nm excitation (# L34957, Life Technologies, 1:500) and APC anti-human CD71 Antibody (#334107, BioLegend, 1:75) for human cell lines and alternatively APC anti-mouse CD71 Antibody (#113820, BioLegend, 1:75) for the murine cell line, for 30 minutes at RT in the dark. The cells were washed with PBS and fixed for 30 minutes in the dark at 4ºC with 2% PFA. After fixation, cells were washed twice with PBS + 2% FBS and then resuspended in 200 μL of FACS buffer. The mixed cell suspensions were then analyzed by flow cytometry using the BD LSRFortessa cytometer with appropriate excitation and detection channels: LIVE/DEAD (405 nm excitation laser), and for CD71 (640 nm excitation laser). Flow cytometry data were analyzed using FlowJo version 10.6.1.

### 2.8 POs uptake studies by flow cytometry

The cells were seeded in 6-well plates at a density of 20 x 10^4^ cells per well and left to grow for 48h. They were then incubated with empty POs labeled with Cy7 dyes at a density of 2.8 x 10^10^ POs per milliliter for 30 min and 2h timepoints. After the incubation, cells were washed with PBS and detached using trypsin. Cells were washed with PBS+2% FBS for 3 min at 1300 rpm and incubated with LIVE/DEAD® Fixable Aqua Dead Cell Stain Kit, for 405 nm excitation (# L34957, Life Technologies, 1:500) to discern live and dead cells. This staining lasted for 30 minutes at RT and in the dark. Afterwards, the cells were washed with 100 μL of PBS and centrifuged at 1300 rpm for 3 min at RT and the supernatant discarded. Next, the cells were fixed by being incubated for 30 min in the dark and at 4ºC with PFA 2%. After incubation, the cells were washed twice with PBS containing 2% FBS and then resuspended in 200 μL of FACS buffer. Finally, the mixed cell suspensions were analyzed by flow cytometry using the BD LSRFortessa cytometer, at the Flow Cytometry Platform of GIMM, with appropriate excitation and detection channels: 405nm excitation laser with 475 LP filter for LIVE/DEAD and 640nm excitation laser with a 750 LP filter for POs detection. Flow cytometry analysis was done using FlowJo version 10.6.1.

### 2.9 Cell viability studies

CRC and HIEC cells were seeded in a 96-well plate at a density of 10^4^ cells per well. After 48 hours the culture medium was replaced with fresh media containing either free DOX or DOX/0%T7pep-POs (DOX-pristine) and DOX/0.25%-T7pep POs, achieving a final DOX concentration of 4.6 μM. Control groups containing only media were included as blanks. Cells were treated for 24 and 48 hours, after which cell viability was assessed using the CyQUANT XTT Cell Viability assay (#X12223, Thermo Fisher Scientific), according to the manufacturer’s instructions. Mitochondrial viability, used as an indirect measure of cell viability, was quantified at both time points. Cytotoxic effects were evaluated by measuring absorbance at 450nm, with a secondary measurement at 660nm to account for autofluorescence, using a TECAN Spark microplate reader.

### 2.10 In vivo studies

C57BL/6J mice were bred, maintained, and used at the Rodent Facility of GIMM under pathogen-free conditions. Animal experimentation was approved by the ORBEA-GiMM (GiMM’s Animal Welfare Body) and DGAV (the Portuguese National Authority for Animal Health) in accordance with Portuguese regulations. To assess the anti-tumor efficacy of DOX-POs in vivo, 7.5x10^4^ MC38 cells were injected into the flank of each mouse (N=21). The animals were then randomly assigned to four treatment groups: PBS (vehicle), free-DOX, DOX/ 0% T7pep-POs, and DOX/0,25%T7pep-POs. After 12 days post-tumor inoculation, mice received intravenous injections of DOX at maximum dose of 2.5 mg/Kg, administered every two days. Tumor growth was monitored, and tumor size was measured using a caliper (#PTR006777, PTROBOTICS). Tumor volume was calculated using the following formula:

. Tumor growth was monitored, and size measured using a caliper (#PTR006777, PTROBOTICS). Tumor volume was obtained according to the following formula:

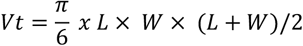

Animals were sacrificed when tumors reached 400 mm^3^ or when tumors became ulcerated. Euthanasia was performed via transcardiac perfusion using 20mL of PBS followed by 20mL of 4% Formaldehyde (#F8775, Sigma-Aldrich). Blood was collected before previously the perfusion procedure. The following tissues were harvested: heart, liver, kidneys, spleen, lymph nodes and tumor, washed in PBS and preserved in 4% PFA overnight and kept in 30%sucrose for subsequent studies on tissue cytotoxicity.

### 2.11 H&E and immunohistochemistry analysis of treated tissues

All fixed organs and tumors collected from the mice were sent to the histology department for processing. Tissues were processed using an overnight protocol in a Tissue HistoCore Pearl (Leica) and embedded in paraffin (#39602012, Leica). Sections of 3μm thickness were obtained using a microtome (Minot Microtome Leica RM2245). Tumor and organ sections were stained with hematoxylin (#0506004E, bio-optica) and eosin (#110132-1L, Sigma-Aldrich) (H&E) for morphological examination. For immunohistochemistry, tumor sections were subjected to heat-induced epitope retrieval, followed by inactivation of endogenous peroxidase using 3% hydrogen peroxide (H_2_O_2_) in methanol for 30 minutes at room temperature. Samples were then blocked with Dako Protein Block for 40 minutes at room temperature, incubated with Caspase 3 primary antibody (#9662S, Cell Signaling Technology, 1:300) (diluted in Dako diluent) for 1h, and washed. Following that, the sections were incubated with the secondary antibody EnVisionTM Flex conjugated with horsedish peroxidase (HRP) (α-rabbit) (#K4010, Dako). After further washing, the sections were incubated with diaminobenzidine (DAB) for 3 minutes, counterstained with Harris Haematoxylin for 10 minutes, and washed again. Both stainings were performed by the Histopathology Facility at GIMM. Finally, the sections were mounted, and images were captured using the Zeiss Axioscan 7 microscope at the Bioimaging Platform of GIMM.

### 2.12 Statistical analysis

Graphs and statistical analysis were performed using GraphPad Prism version 8.4.2. Statistical significance is reported as follows: p≤ 0.05 (*), p ≤ 0.01 (**), p≤ 0.001(***) and p≤ 0.0001 (****). A One-way ANOVA paired with Dunnet’s multiple comparisons test was used to compare the mean fluorescence intensity (MFI) of cellular uptake of POs with different molar concentrations of T7 peptide between two CRC cell lines. Two-way ANOVA with Dunnett’s multiple comparisons test was used to compare cellular uptake of POs with different molar concentrations of T7 peptide in each cell line, as well as for the cell viability assays. One-way ANOVAS was applied to compare linear regressions of tumor growth. The Gehan-Breslow-Wilcoxon test was used to compare survival between the tested mouse groups.

## 3 Results

### 3.1 Design and characterization of empty T7pep-POs targeting transferrin receptor

Considering our aim to use POs as drug delivery systems targeting specific receptors, such as TfR1, it is crucial to analyze their size [17]. We first designed and prepared various formulations of POs targeting TfR1 by decorating their surface with different molar ratios of T7 peptide (HAIYPRH) using the solvent displacement method. The POs were then characterized by DLS to evaluate their size (nm), homogeneity, and quality. Light scattering analysis revealed that functionalization with different molar ratios of T7pep on the surface of PEG-PLA POs did not alter their size (Figure 1A & B), with an average hydrodynamic diameter (D_h_) of 88.8 nm (Table S1). TEM analysis of T7pep-POs confirmed the formation of spherical vesicles, with Dh size consistent with the DLS measurements (Figure 1C and Figure S1). All PO formulations exhibited a narrow size distribution, as indicated by a polydispersity index (PDI) of less than 0.3, suggesting good homogeneity and average zeta potential of - 0.4mV, indicating an inert surface charge. Therefore, functionalization with increasing molar ratios of T7pep on the surface of POs did not affect the size, resulting in a monodispersed formulation with no evidence of aggregation, making the T7pep-POs suitable for further use in biological samples.

**Figure 1.**
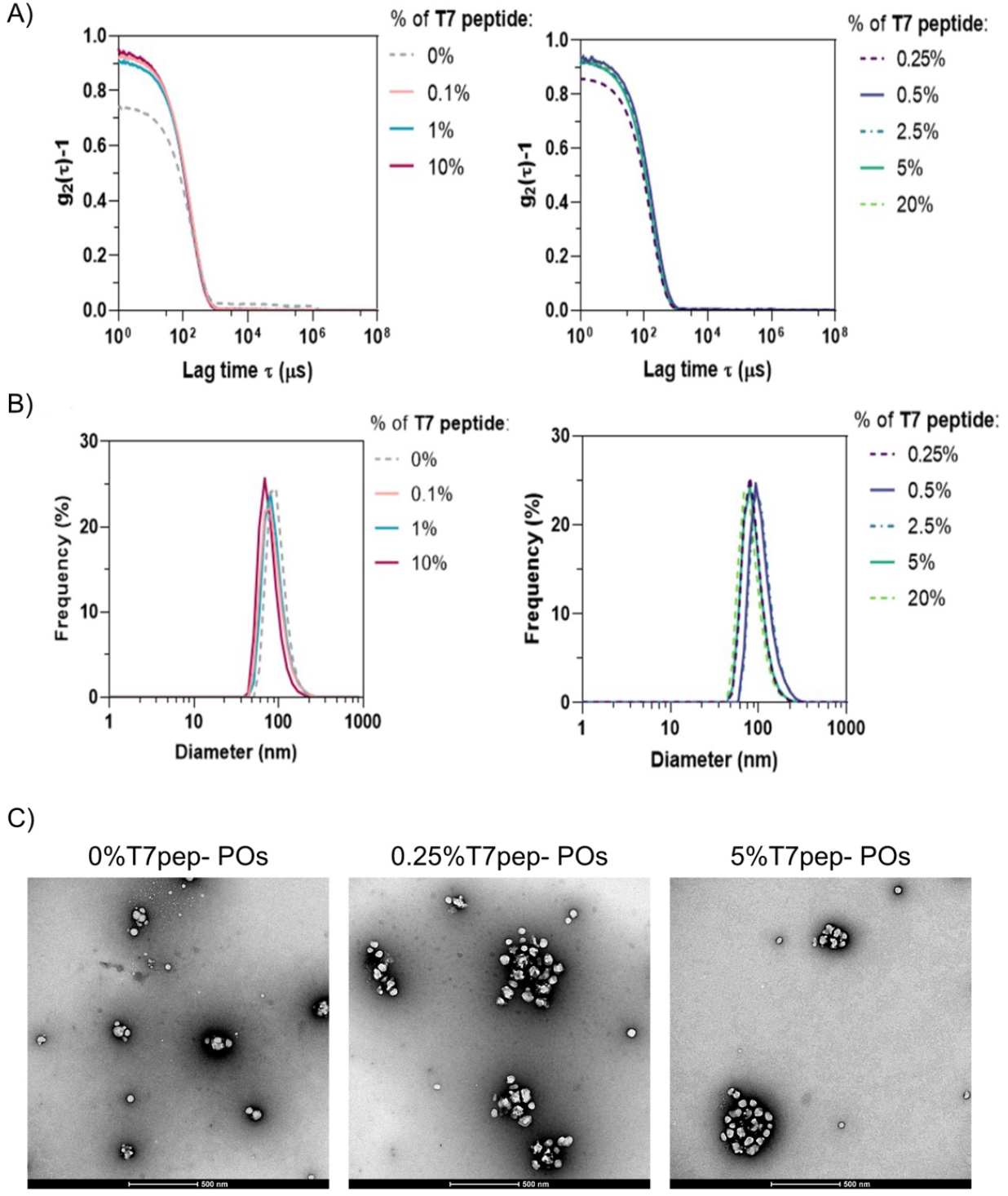
Characterization of T7pep-POs by DLS and TEM. **A)** Graphs showing the DLS autocorrelation function g_2_(*t*)-1 for different molar percentages of T7pep-POs **B)** DLS number-averaged distribution calculated from the correlation functions in **A**. The intensity-averaged distribution confirms the absence of aggregate formation. All values represent three independent experiments (N=3). **C)** TEM images of three formulations: pristine (0%), 0.25% and 5% T7pep-POs. Scale bar= 500 nm.

### 3.2 Uptake of empty POs in healthy and CRC cell lines

To characterize TfR1 expression in our cell models, we first assessed its cellular distribution through immunofluorescence (Figure 2A), confirming the presence of TfR1 on the cell surface in all cell lines tested. To quantitatively evaluate TfR1 levels, we performed flow cytometry (Figure S2A), revealing a significantly higher percentage of TfR1-positive cells in CRC cell lines compared to hIECs (p < 0.0001). Additionally, analysis of the median fluorescence intensity (MFI) (Figure 2B) demonstrated that the murine CRC cell line expressed notably higher levels of TfR1 than the human CRC and healthy cells (p < 0.0001). Therefore, to assess the specificity of our POs for targeting TfR1 in our cell models, we first optimized the incubation conditions to avoid saturation. HCT116 cells were incubated for 2 hours with varying number of 0% and 0.25%-T7pep-POs and cellular uptake was assessed using flow cytometry (Figure 2C & Figure S2B). The results identified an optimal number of 2.8 x 10^10^ POs, which minimized nonspecific binding of pristine POs while still allowing the binding of 0.25%-T7pep-POs. Next, we repeated the cellular uptake assay with all PO formulations in HCT116, HT29 (Figure 2D & E), left and right, respectively), MC38 and hIECs (Figure S2C). In the HCT116 cells, only the 0.25% and 1%T7pep-POs exhibited nearly 100% uptake at 2hours, with both formulations showing significant differences compared to 0%T7pep-POs (p < 0.0001) (Figure 2D). A similar trend was observed in the HT29 cell line (Figure 2E). In the MC38 and hIEC cell lines, after 2 hours, the 0.25% and 1%T7pep-POs also displayed higher specific uptake compared to the other formulations (Figure S2C). Notably, cellular uptake was lower in the healthy hIECs cells compared to the CRC cells (Figure S2C). These findings highlight the differential expression of TfR1 across cell types and confirm the specificity of T7pep-POs for CRC cells, with significantly higher uptake in CRC cell lines compared to healthy hIECs. Based on these results, the optimal formulation of 0.25%T7pep-POs and the corresponding number of POs (2.8 x 10 ^10^) were selected for further studies.

**Figure 2.**
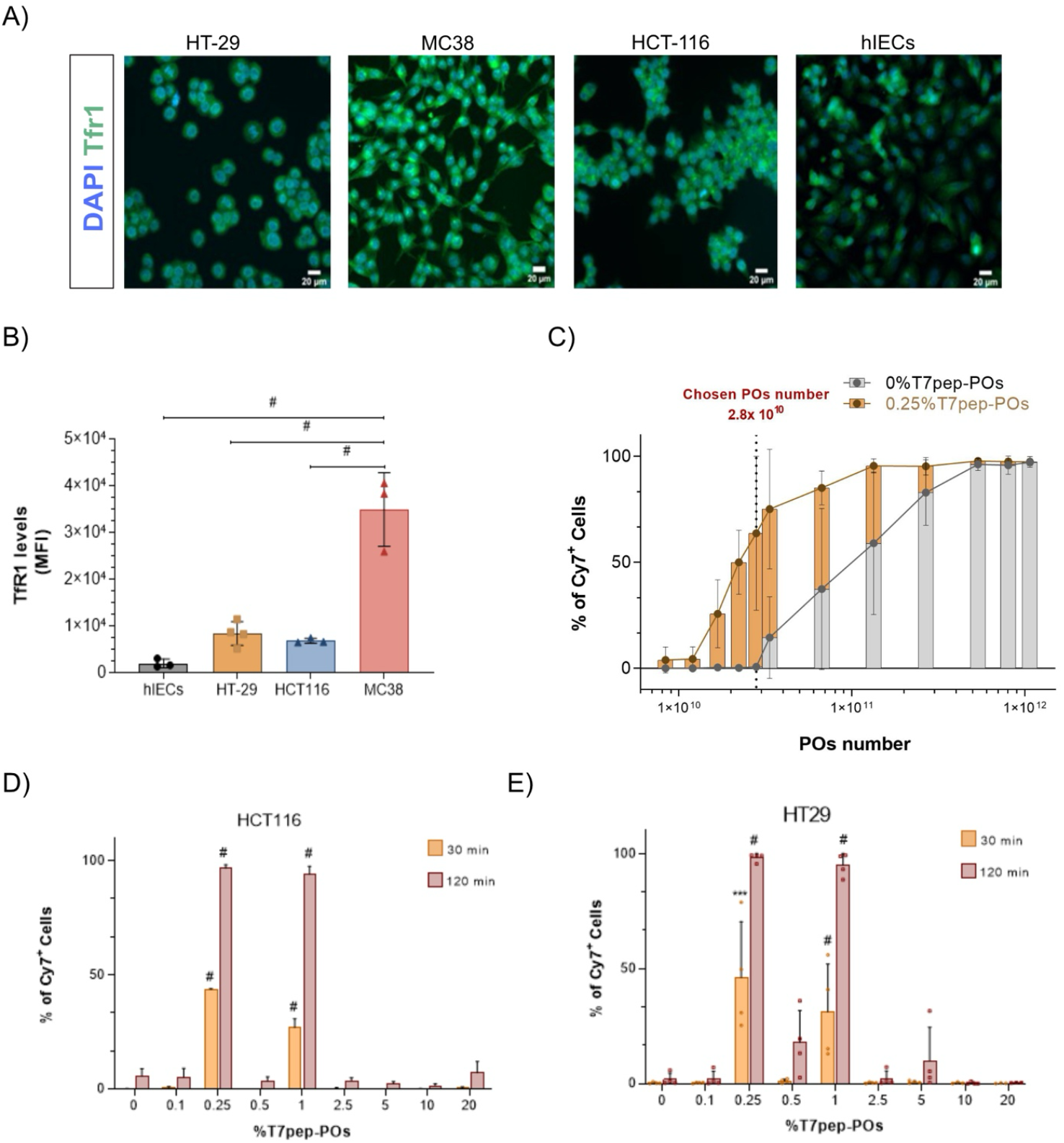
TfR1 expression and cellular uptake of T7pep-POs in CRC and healthy hIEC cells. **A)** Cellular distribution of TfR1 in HT-29, HCT116, MC38 CRC cells and hIECs by immunofluorescence. Images were captured using a Zeiss Cell observer microscope equipped with a Photometrics Prime BSI Express sCMOS camera. Images acquired using an EC Plan-NeoFluar objective with a 40X magnification. **B)** Expression levels of TfR1 based on median fluorescence intensity (MFI) measured by flow cytometry analysis in CRC and healthy cell lines. **C)** Percentage of cellular uptake of different number of 0%T7pep-POs (Pristine) after 2 hours incubation(N=4). **D)** Cellular uptake of different molar ratios of T7-pep-POs (NPOs=2.8 x 10^10^) for HCT116 (N=4) and E) HT-29 (N=4). Statistical analysis using a two-way ANOVA, statistically significant when p≤ 0.001(***) and p≤ 0.0001 (#).

### 3.3 Effect of DOX-encapsulated TfR1-targeted POs on CRC cells in vitro

Based on the results of the cellular uptake assay, we selected the 0.25%T7pep-POs for encapsulating DOX, as it exhibited the highest cellular uptake in CRC cells (Figure 2D & E). The DOX-loaded POs were characterized by DLS to assess their size, homogeneity, and quality. Our data showed that the size of 0.25%T7pep-POs with DOX was 74.83 nm (Figure 3A, B & C and Table S2), indicating that encapsulation did not significantly affect the size compared to non-encapsulated POs (Figure 3C). However, the DOX/0%T7pep-POs were similar (78.24 nm) as the empty POs (Figure 3C). TEM analysis confirmed the formation of spherical vesicles for DOX/T7pep-POs, with sizes consistent with the DLS measurements. Additionally, the encapsulation amount of DOX in 0%T7pep-POs was 42.86 μg/mL, while in the 0.25%T7pep-POs was 46.9 μg/mL, corresponding to encapsulation efficiencies of 2.14 and 2.35%, respectively (Table S2). Following the successful encapsulation of DOX in both 0%T7pep-POs and 0.25%T7-POs, we assessed their cytotoxic effect in CRC cells and compared them to free DOX *in vitro*. Cells were incubated with 4.6 μM of DOX, either as a free drug or encapsulated within POs, for 24 and 48 hours (Figure 3D). Among the tested cell lines, HCT116 (Figure 3D, top left) showed the highest sensitivity, with significantly reduced viability at 48 hours, compared to untreated cells (p < 0.0001). In HT-29 cells (Figure 3D, top right), the DOX/0%T7pep-POs unexpectedly exhibited the strongest cytotoxic effect. However, the DOX/0.25%T7pep-POs showed a more pronounced effect (p < 0.0001) than free DOX (p = 0.0002) at 48 hours. In MC38 cells (Figure 3D, bottom left), free DOX has the strongest effect at 24 hours (p = 0.0004), but after 48 hours, DOX/0.25%T7pep-POs exhibited a greater cytotoxicity than other formulations (p=0,0099) and even surpassed free DOX (p = 0.0175). For the healthy cell line (Figure 3D, bottom right), minimal cytotoxic effects were observed at 24 hours, with the DOX/0.25%T7pep-POs showing no significant toxicity (p = 0.4762). However, at 48 hours, their cytotoxicity increased, becoming comparable to free DOX (p < 0.0001). These findings suggest that DOX/0.25%T7-pep-POs effectively deliver DOX to CRC cells, enhancing cytotoxicity while potentially reducing toxicity to healthy cells (hIECs).

**Figure 3.**
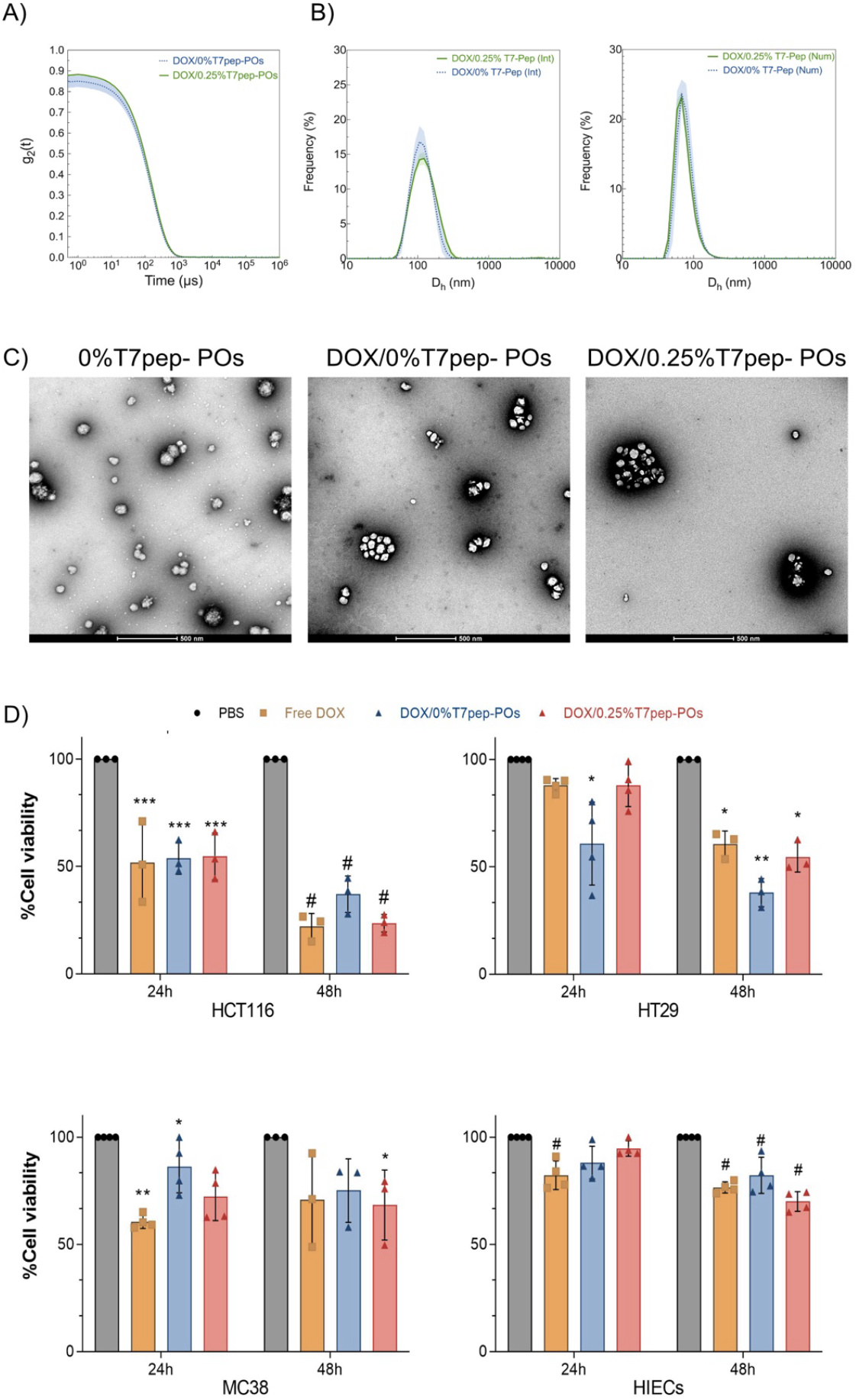
Cytotoxic effects of DOX-loaded T7pep-POs on CRC and healthy cells. A) Graphs showing the DLS autocorrelation function g^2^(*t*)-1 for DOX/0%T7pep-POs and DOX/0.25%T7pep-POs. **B)** Intensity-averaged (left) and number-averaged (right) DLS size distribution (N=4). **C)** TEM images of empty 0%T7pep-POs and DOX-loaded T7pep-POs. Scale bar= 500 nm. **D)** Cells treated with 4.6 μM of free DOX, DOX/0%T7pep-POs, or DOX/0.25%T7pep-POs for 24 and 48 hours. Cell viability was assessed using the XTT in HTC116 **(Top right)**, HT29 **(Top left)**, MC38 **(bottom left)** and hIECs **(bottom right)**. Statistical analysis was performed using two-way ANOVA with significance reported as p< 0.05 (*), p < 0.01 (**), p< 0.001(***) and p< 0.0001 (#).

### 3.4 Evaluation of DOX-loaded T7pep-POs efficacy in in vivo CRC model

To test the effectiveness of the DOX-loaded T7pep-POs in a murine CRC model, we conducted *in vivo* experiments. MC38 cells were injected into C57BL/6J male mice, and after 12 days post-injection when tumor was already visible (on average, 22.1 mm^3^), we started the treatments. Mice were assigned to four treatment groups: PBS (control), free DOX, DOX/0%T7pep-POs and DOX/0.25%T7pep-POs (Figure 4A). Survival analysis (Figure 4B) demonstrated that DOX/0.25%T7pep-POs significantly prolonged survival compared to both free DOX treatment group (p = 0.0082) and PBS control group (p = 0.0216). Unexpectedly, mice receiving free DOX exhibited the lowest survival rate among all groups. Tumor growth was monitored throughout the study, and linear regression analysis was performed to evaluate growth trends (Figure 4C). Both DOX/T7-POs formulations reduced tumor progression, with DOX/0.25%T7pep-POs showing the most pronounced effect. This formulation significantly inhibited tumor growth compared to PBS (p = 0.0306) and free DOX (p = 0.0116), suggesting its enhanced therapeutic efficacy. To further evaluate the treatment’s impact, tumor and tissue samples were collected and analyzed by H&E staining. Histological analysis confirmed that DOX/0.25%T7pep-POs induced extensive necrosis within tumor tissues (Figure 4D) while sparing healthy tissues, with no detectable damage observed (Figure S3), reinforcing its potential as a targeted and effective CRC therapy.

**Figure 4.**
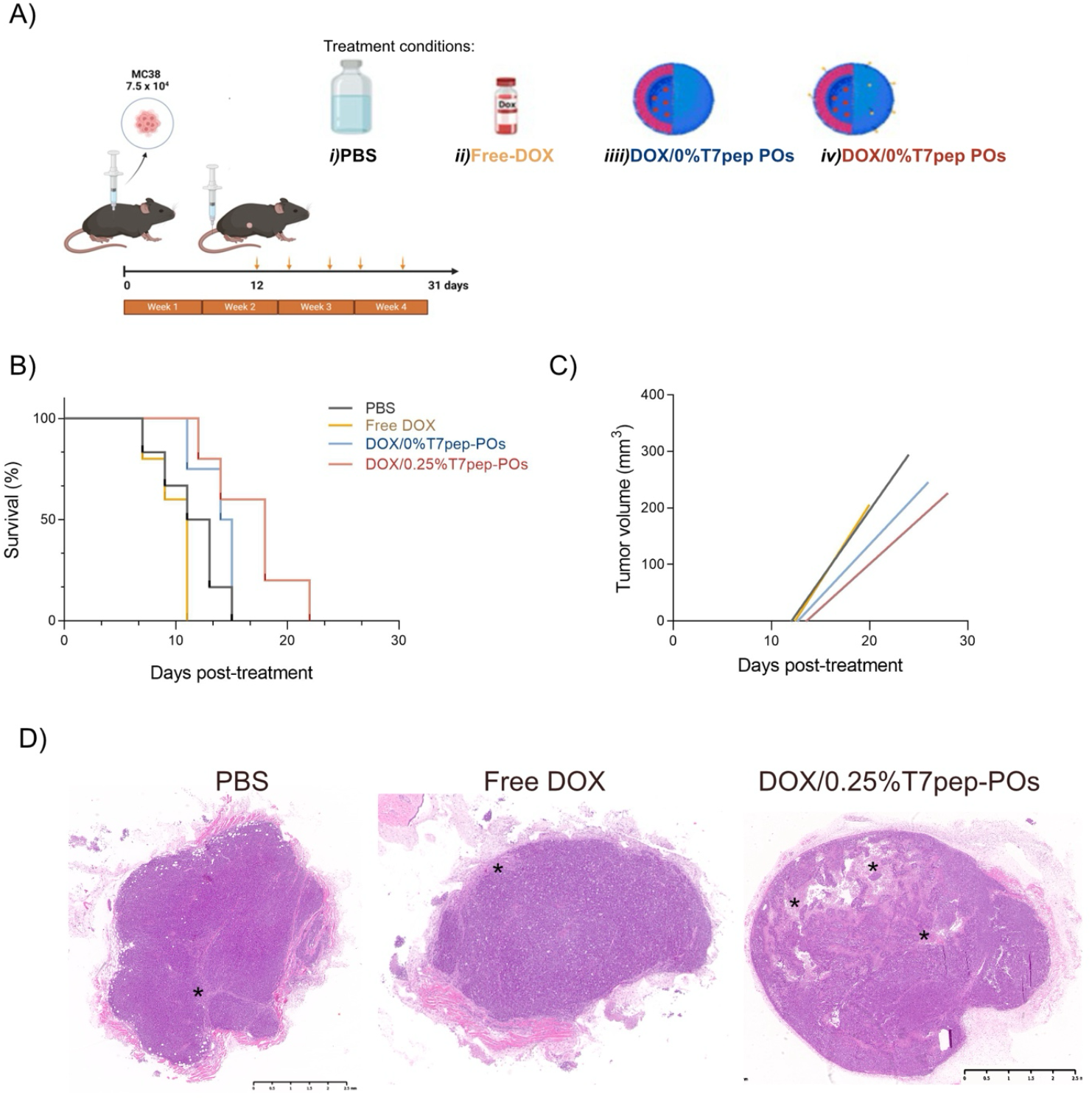
Effect of DOX-loaded T7pep-POs in vivo. **A)** Experimental in vivo design scheme. **B)** Survival curves showing the duration of survival from the first day of treatment. Mice were divided into four experimental groups: PBS (N = 6), free DOX (N = 5), DOX/0%T7pep-POs (N = 4); DOX/0.25%T7pep-POs (N = 5). **(C)** Linear regression analysis representing tumor growth across all treatment groups. Tumor growth was analyzed using linear regression followed by a one-way ANOVA with a multiple comparison. Survival analysis was performed using a Log-rank (Mantel-Cox) test. **D)** Hematoxylin and eosin (H&E) staining of tumor sections representative of the PBS, free DOX and DOX/0.25% T7pep-POs treatment groups. * Indicates necrotic areas.

## 4 Discussion

CRC is a highly heterogenous malignancy characterized by significant innate and acquired resistance to treatments, making it challenging to cure and affecting millions of patients worldwide [1,2]. Standard therapeutic approaches, primarily based on chemotherapeutic agents, can extend a patient survival, but often induce severe side effects, including neuro-, cardio-, hepato-and nephrotoxicity due to their lack of specificity [4,7]. This highlights the urgent need for more targeted treatments that are not only effectively eliminate cancer cells but also minimize harm to healthy tissues, ultimately enhancing patient outcomes and quality of life. One promising target for such therapies is TfR1, overexpressed in tumor cells, particularly in CRC [13]. This makes TfR1 an ideal target for testing NP-based therapies, such as POs which can work as drug delivery systems [14,15,18]. In our study, we demonstrated that TfR1-targeted POs delivered DOX intracellularly and induced cell death of tumor cells. We first generated different TfR1-targeting POs and confirmed TfR1 expression in our cell models before performing cellular uptake studies. Since at 0.1 mg/mL concentration of POs, we observed a saturation of the receptors in our system, prompted us to refine our approach by testing various PO concentrations in the HCT116 cell line at 2hours. This included both the pristine POs and the more specific 0.25%T7pep-POs. Upon optimizing the assay conditions, we observed notable differences in uptake. Notably, cells treated with 0.25% and 1%T7pep-POs exhibited significantly higher uptake, indicating enhanced specificity. These results surpassed previously reported NP formulations used in CRC research, even at lower NP concentrations. For instance, both formulations achieved greater cellular uptake than the T7-liposomes described by Wang & Sun, even when tested at reduced concentrations [19]. Regarding healthy cells, there are, to our knowledge, no published studies on TfR1-mediated uptake. However, it is well-established that healthy cells express lower TfR1 levels than CRC cells, which likely explains the lower affinity and reduced PO uptake observed in these cells [13,20]. After identifying the optimal T7pep-PO candidate with superior affinity and barrier-crossing properties, we successfully encapsulated DOX into both pristine and 0.25%T7pep-POs. Subsequent cell viability studies yielded promising results, particularly considering the relatively low DOX concentration used (4.6 μM). Wei et al showed that PEG-poly (trimethylene carbonate-co-dithiolane trimethylene carbonate) POs functionalized with the TfR binding peptide (CGGGHKYLRW) and encapsulated DOX in HCT116 reduced 45% of cell viability after 72 hours. While our T7pep-POs at 48h were already much more effective with 20-30% of viable cells [21]. Similarly, when compared to T7-liposomes, our POs exhibited higher cytotoxic against CRC cell lines at 48 hours, despite using different therapeutic agents. At equivalent therapeutic concentrations, our POs achieved lower cell viability than the liposomes, which maintained 75% cellular viability [19]. These differences may stem from variations in the mechanisms of action between DOX and other therapeutic agents, as well as the superior stability of POs compared to liposomes. Unlike liposomes, POs can evade the immune system and maintain structural integrity under physiological conditions [14,15,19,22]. Interestingly, the pristine formulation also had a strong cytotoxic effect in CRC cell lines, even in absence of the T7pep ligand. This finding aligns with previous studies demonstrating that pristine formulations can reduce cell viability proportionally to the amount of encapsulated therapeutic agent, suggesting this is a common henomenon [19,21]. In in vivo assays, DOX/0.25% T7pep-POs effectively suppressed tumor growth, reducing the average tumor size to 300 mm^3^ by day 18 post-treatment, consistent with findings by Alibolandi et al. However, survival outcomes differed significantly. Mice treated with DOX/0.25%T7pep-POs in our study survived for 21 days after treatment initiation, while Alibolandi et al., reported survival extending up to 40 days [23]. This discrepancy may be attributed to differences in euthanasia criteria and targeting, as their study used POs to target the integrins (αvβ3). In our study, mice were euthanized when tumors reached 400 mm^3^ or ulcerated in the case of free DOX and PBS groups. In contrast, the other work, they euthanized mice only when tumors reached 2000 mm^3^ or impaired feeding [23]. Additionally, the use of different cell lines, sexes, and animal models, C26 cells (BALB/c, female) and MC38 cells (C57BL/6, male) in our study, may have contributed to the observed variations. [23]. Histopathological analysis further revealed minimal necrosis in the tumor tissue from PBS and free DOX-treated groups compared to the DOX/0.25%T7pep-POs group. The limited necrotic effect of free DOX likely reflects its dependency on dose and administration frequency. Wei et al, demonstrated that higher DOX doses (e.g. 4 mg/kg) induce both tumor and healthy tissue necrosis [24–26]. In contrast, the lower DOX concentration used in our study, was 2.5 mg/kg, standing on the lower end of the range reported in the literature (i.e. 2-40 mg/kg), which likely explains the reduced necrotic effect on the tumors and lack of necrotic effect on healthy tissues [24]. Additionally, the absence of healthy organ damage following free DOX administration further supports the notion that higher dosages are required for such adverse effects [21,24]. Despite the relatively low DOX concentration, the DOX/0.25%T7pep-POs demonstrated significant anti-tumoral activity while sparing healthy tissues. These findings underscore the potential of T7pep-functionalized POs as a promising therapeutic strategy for CRC treatment, offering an effective and less toxic alternative to conventional chemotherapy.

## 5 Conclusions

Although TfR1 has been a key focus in cancer therapy research for decades, particularly in brain-related malignancies, its exploration in the context of CRC remains relatively limited. Our findings strongly reinforce the potential of TfR1 as a viable target for drug delivery systems in CRC, highlighting the promise of our TfR1-targeting POs in advancing more precise and effective therapeutic strategies. The observed variations in outcomes between studies can be attributed to differences in euthanasia criteria, cell lines, animal models, and DOX dosing regimens. In this study, the lower DOX concentration (2.5 mg/kg) likely explains the reduced necrotic effect on tumors and the absence of healthy tissue damage compared to higher doses reported in the literature. Despite the lower dose, DOX/0.25% T7pep-POs demonstrated significant anti-tumoral activity while sparing healthy tissues. These findings highlight the potential of T7pep-functionalized POs as an effective and less toxic therapeutic strategy for CRC treatment, offering a safer alternative to conventional chemotherapy. Notably, the 0.25% T7pep-POs exhibited remarkable specificity in targeting CRC cells through T7 peptide functionalization, further emphasizing its promise as an innovative treatment approach for this challenging malignancy.

## Acknowledgments

We would like to thank the Flow cytometry, Histopathology, Bioimaging (funded by PPBI-POCI-01-0145-FEDER-022122) and Rodent platforms of the GIMM for their technical support. We would also like to acknowledge Michael Hall from the Electron Microscopy Facility for technical expertise, sample processing and imaging.

## Author contributions

**Ariana Pina:** Writing-review & Editing, Writing-original draft, Visualization, Validation, Methodology, Investigation, Formal analysis, Data curation, Conceptualization. **Elisa Mastrantuono**: Data curation, investigation, methodology, validation, formal analysis. **Marta Silva:** Data curation, investigation, methodology, validation. **Valentino Barbieri:** Writing-original draft, Methodology, Investigation, Formal analysis. **José Muñoz-López:** Data curation, investigation, methodology, validation, formal analysis. **Giuseppe Battaglia:** Writing-review & Editing, Project administration, Funding acquisition **Luis Graça:** Writing-review & Editing, Project administration, Funding acquisition **Diana Matias:** Writing-review & Editing, Writing-original draft, Visualization, Validation, Supervision, Resources, Project administration, Methodology, Investigation, Funding acquisition, Formal analysis, Data curation, Conceptualization.

