## Supplementary material for "Transferrin Receptor 1-targeted polymersomes therapy for Colorectal Cancer": Figure S1

**Supplemental Figure 1**

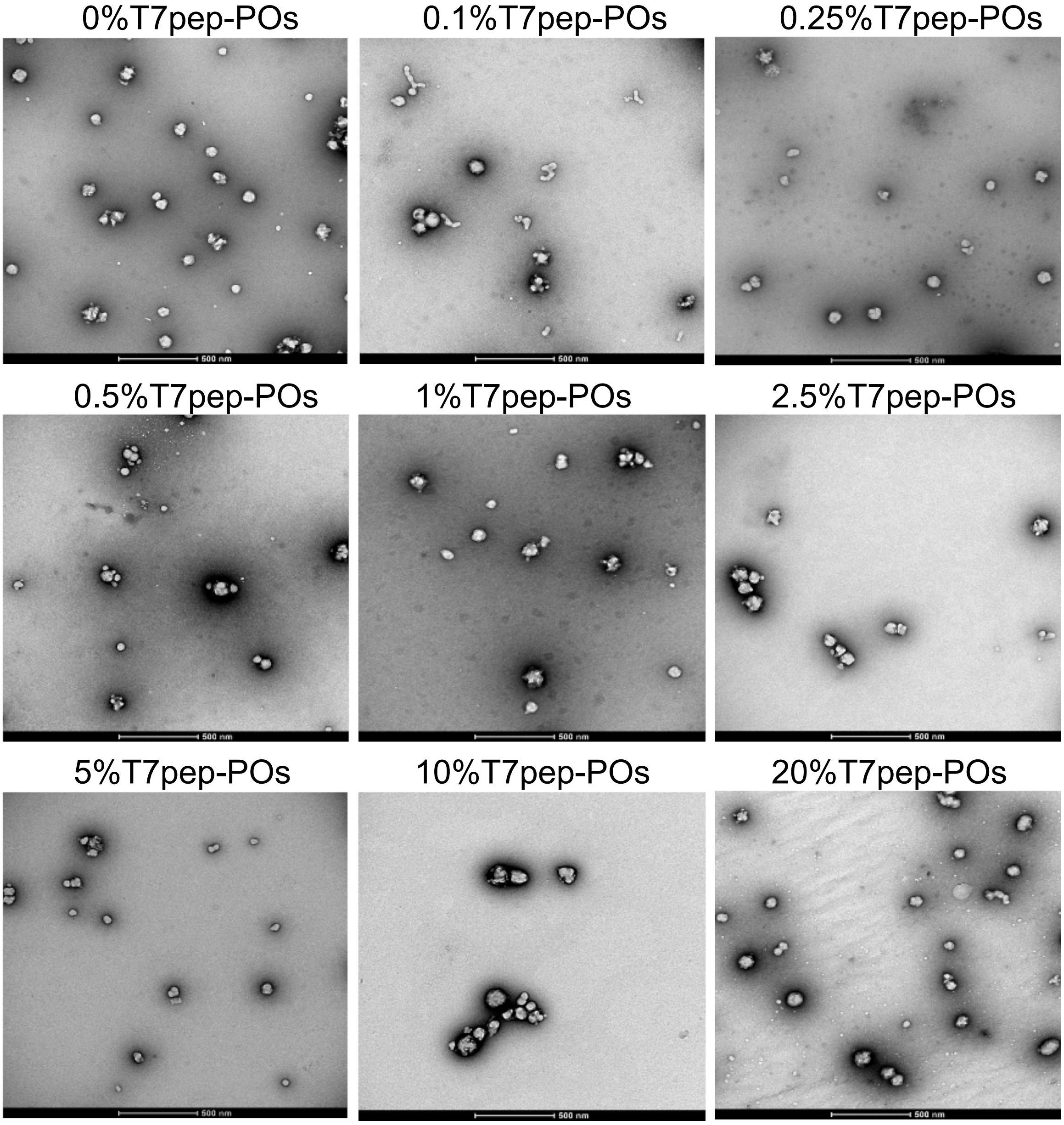


***Figure S1- TEM images of different T7pep-POs.*** TEM images of the 9 POs formulations functionalized with T7 peptide (0% to 20%). The images display the morphology and size distribution of the POs, illustrating the successful incorporation of T7 peptide into the POs and confirming the formation of spherical vesicles. Scale bars represent 500 nm.
