## Supplementary material for "Transferrin Receptor 1-targeted polymersomes therapy for Colorectal Cancer": Figure S2

### Supplemental Figure 2

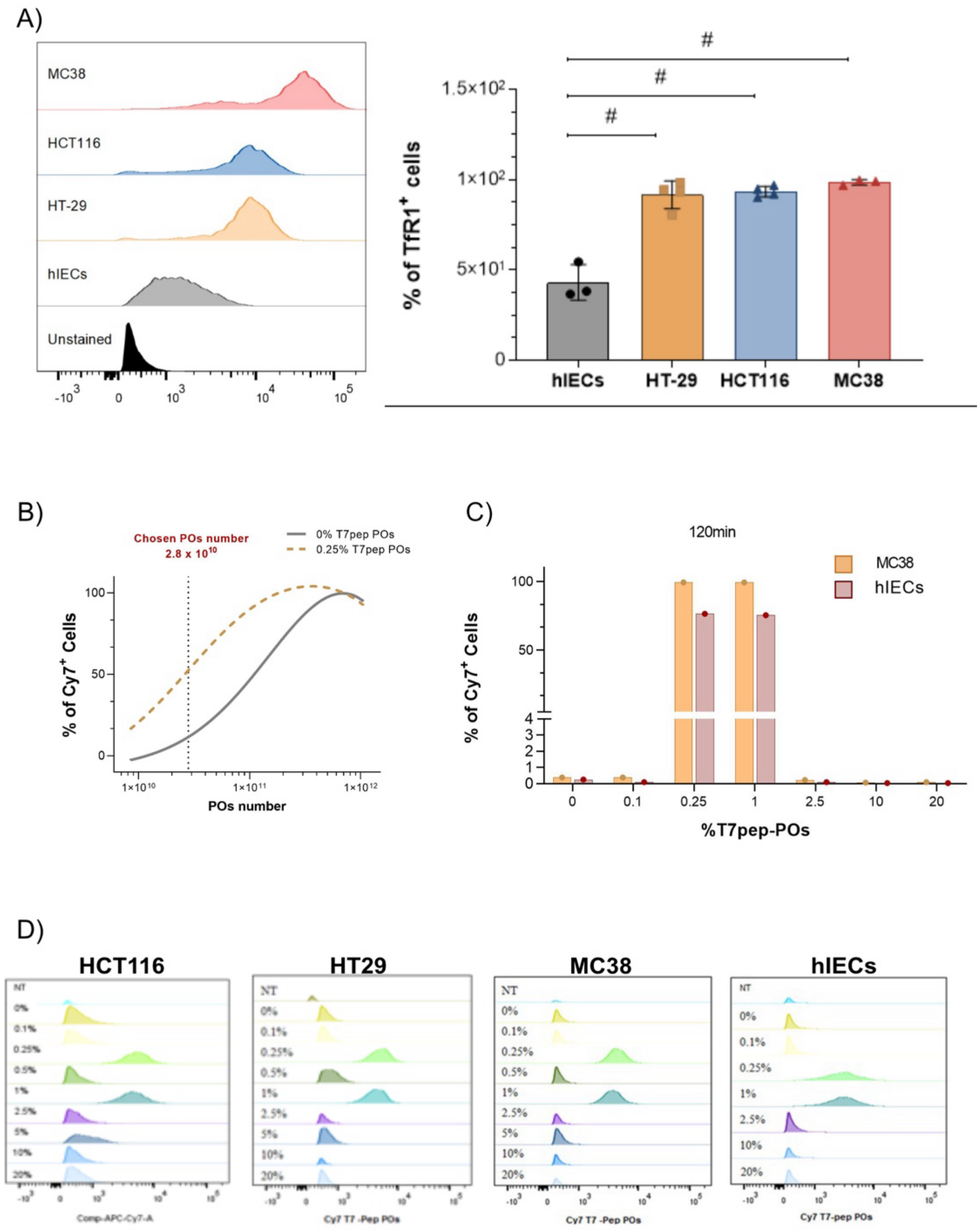


***Figure S2- TfR1 expression and cellular uptake of T7pep-POs in MRC38 and healthy hIEC cells at 2 hours.*** **A)** Histograms representing the expression levels of TfR1 based on MFI distribution (*left*) and the percentage of cells expressing the receptor in CRC and healthy cell lines (*right*). **B)** Optimization of the number of 0% T7pep-POs (Pristine) and 0.25% T7pep-POs in HCT116 cells (N=4). **C)** Cellular uptake of different molar ratios of T7pep-POs (NPOs = 2.8 x 10^10) in MC38 (N=1) and hIEC (N=1) cell lines. **D)** Histogram of cellular uptake of T7pep-POs (NPOs = 2.8 x 10^10) in CRC and healthy cell lines. Statistical analysis was performed using a two-way ANOVA, with statistical significance denoted as p ≤ 0.001 (***) and p ≤ 0.0001(#).
