## Supplementary material for "Transferrin Receptor 1-targeted polymersomes therapy for Colorectal Cancer": Figure S3

### Supplemental Figure 3

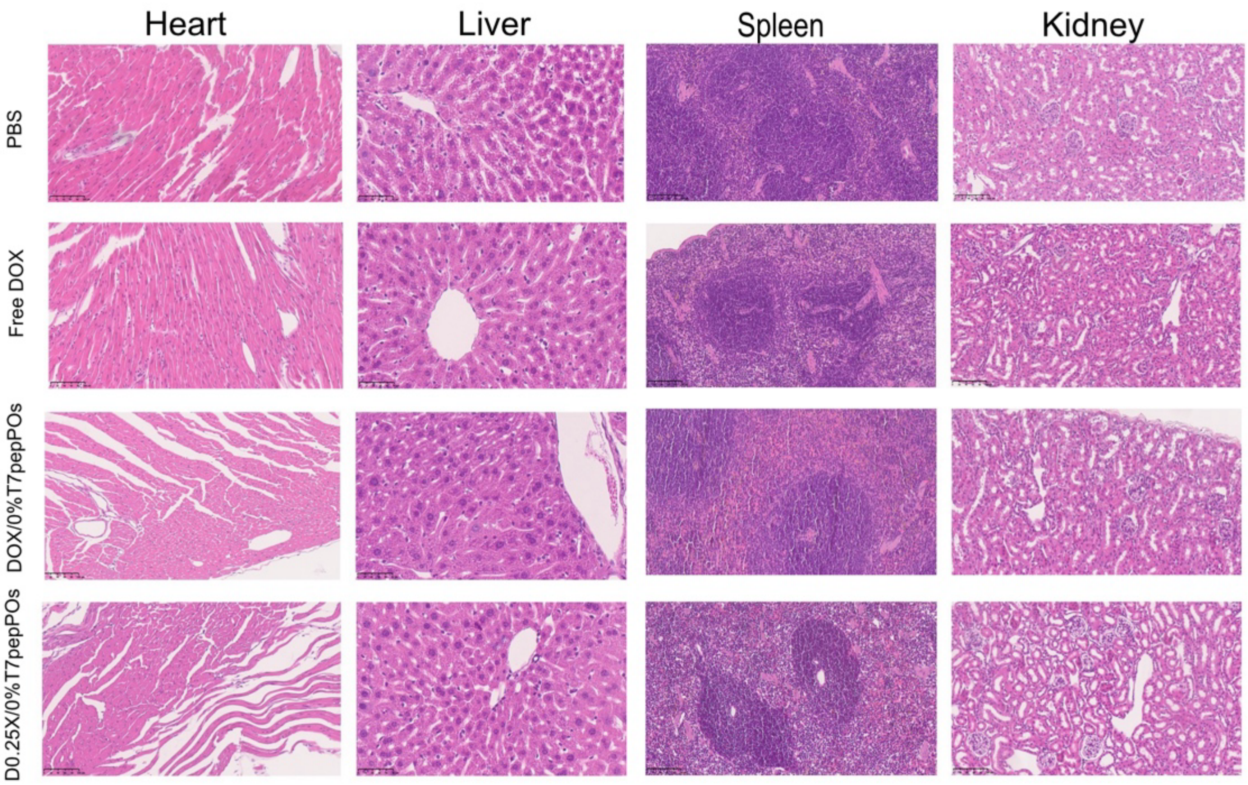


**Figure S3 - Histology of healthy tissues post-treatment.** H&E stainings of tissue (Heart, Liver, Kidneys and Spleen) sections representative of the PBS, free DOX, DOX/0%T7pep-POs and DOX/0.25% T7pep-POs treatment groups. No damage was detected in these tissues.
