## Supplementary material for "Transferrin Receptor 1-targeted polymersomes therapy for Colorectal Cancer": Table S1

### Supplemental Table 1

**Table 1 – Summary data of characterization of POs decorated with % molar ratio of T7 peptide (HAIYPRH) by DLS.**

| % of T7 peptide (HAIYPRH) | Number of ligands | Mean Diameter (nm) | Concentration (mg/mL) |
|---|---|---|---|
| **0%** | 0 | 83.12 | 3.99 |
| **0.1%** | 9 | 91.9 | 5.01 |
| **0.25%** | 29 | 97.46 | 4.08 |
| **0.5%** | 73 | 103.08 | 5.64 |
| **1%** | 78 | 87.96 | 4.5 |
| **2.5%** | 206 | 87.82 | 6.07 |
| **5%** | 524 | 93.92 | 4.47 |
| **10%** | 380 | 67.08 | 3.68 |
| **20%** | 1487 | 87.06 | 4.1 |
