## Supplementary material for "Transferrin Receptor 1-targeted polymersomes therapy for Colorectal Cancer": Table S2

### Supplemental Table 2

**Table 2 – Summary data of characterization and concentration of DOX present in T7pep-POs by DLS.**

| % of T7 peptide (HAIYPRH) | Mean Diameter (nm) | Encapsulated DOX concentration (µg/mL) | Encapsulation efficiency (%) | Zeta Potential (mV) |
|---|---|---|---|---|
| **0%** | 78.24 | 42.86 | 2.14 | -0,588 |
| **0.25%** | 74.83 | 46.9 | 2.35 | -0,171 |
